# The beginnings of light-dependent nuclear relocation in land plants

**DOI:** 10.1101/2024.09.11.611950

**Authors:** Kosei Iwabuchi, Hiroki Yagi, Kenta C. Moriya, Aino Komatsu, Yuuki Sakai, Tomoo Shimada, Ryuichi Nishihama, Takayuki Kohchi, Akiko Harada, Yo-hei Watanabe, Haruko Ueda, Ikuko Hara-Nishimura

## Abstract

The terrestrialization of plants was accompanied by exposure to several environmental stresses. Adaptation to these stresses required numerous changes at the cellular and molecular level. One such adaptation in the leaves of *Arabidopsis thaliana* is the movement of cell nuclei to avoid UV damage. In the dark, the nuclei locate to the bottom walls of leaf cells to distance genetic material from external stresses, but in response to intense blue light (an indication of the presence of UV), they move to the side walls to escape UV-induced DNA damage^1^. The movement is driven by the photoreceptor phototropin and the actin cytoskeleton^2^. However, how this protective mechanism evolved in land plants remains unclear. Here, we show that in the liverwort *Marchantia polymorpha*, nuclei show a similar, but less stable movement in response to intense blue light. In the dark, *M. polymorpha* positioned nuclei on the upper walls of epidermal cells in young thalli, but in response to intense blue light, the nuclei immediately moved to the side walls, similar to *A. thaliana*. However, the movement was transient and the nuclei returned to the upper walls through both the actin and microtubule cytoskeletons.

Unlike *A. thaliana*, *M. polymorpha* responded to prolonged (> 1 day) exposure to low light by moving nuclei from the upper to the side walls through both the actin and microtubule cytoskeletons and two photoreceptors (phototropin and phytochrome). However, no light-dependent nuclear relocation was observed in charophyte algae, suggesting that light-dependent nuclear relocation was initially established in the common ancestor of land plants as a result of terrestrialization and then diverged during land plant evolution.

---

We investigated whether nuclear positioning occurs in *M. polymorpha* by focusing on epidermal cells near the apical notches of the young thalli (called gemmalings) (Extended Data Fig. 1). In the dark, *M. polymorpha* positioned the nuclei on the upper walls of the cells (Fig. 1a, arrowheads), in contrast to their positioning on the bottom walls of leaf cells in *A. thaliana*^3^. For quantitative analysis, nuclei were categorized into three positions: upper walls, side walls, and bottom walls (Fig. 1b). More than 90% of the nuclei were positioned on the upper walls with almost no nuclei positioned on the bottom walls (Fig. 1c, Wild type).

**Fig. 1.**
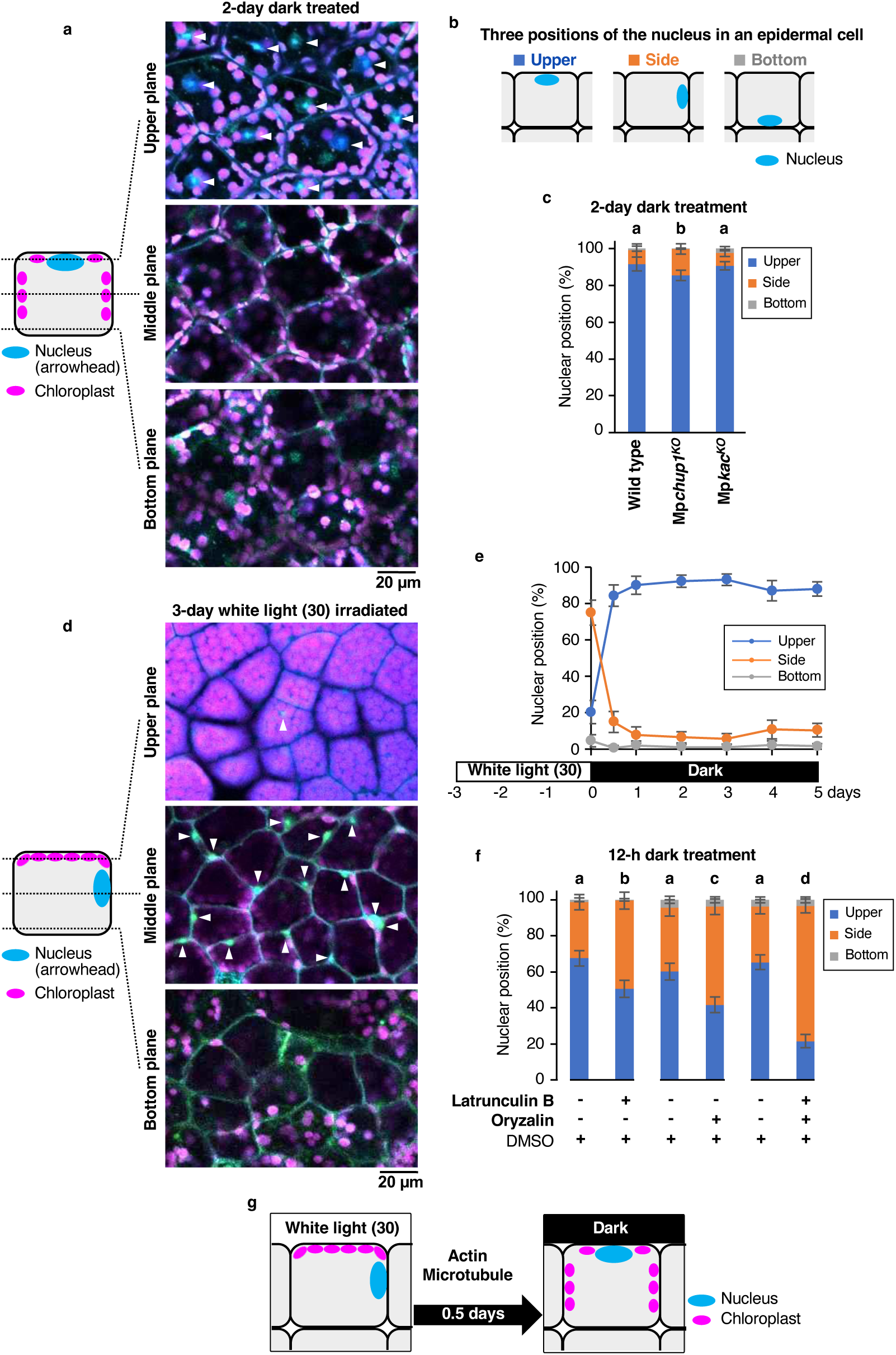
| Dark-induced nuclear relocation depends on two cytoskeletal components in *M. polymorpha*. **a**, Representative images of cells from 2-day-dark-treated gemmalings, showing nuclei stained with DAPI (blue, arrowhead) and chloroplasts (chlorophyll autofluorescence, magenta) located in the upper, middle, and bottom planes of the cells. **b**, Nuclei are categorized into three positions in this study. **c**, Dark-induced nuclear relocation in the wild type, Mp*chup1^KO^*, and Mp*kac^KO^*. Ratios of nuclei at each position are shown. Error bars indicate the 95% CI; n = 205–656 cells, five independent experiments. See Extended Data Fig. 3 for Mp*chup1^ld^*and Mp*kac^ge^*. **d**, Representative images of cells from 3-day-low light-grown gemmalings, showing nuclei stained with DAPI (blue, arrowhead) and chloroplasts (chlorophyll autofluorescence, magenta) located in the upper, middle, and bottom planes of the cells. **e**, Time course of dark-induced nuclear relocation. Ratios of nuclei at each position are shown. Error bars indicate the 95% CI; n = 141–259 cells, five independent experiments. **f**, Effects of cytoskeletal inhibitors on dark-induced nuclear relocation. Ratios of nuclei at each position are shown. Error bars indicate the 95% CI; n = 426–525 cells, five independent experiments. **g**, A mode of dark-induced nuclear relocation in *M. polymorpha* gemmalings. Different letters above the bars indicate statistically significant difference (**c** and **f**).

Unexpectedly, under the artificially prolonged (> 3 days) exposure to continuous low white light (30 µmol m^-2^ s^-1^), most of the nuclei were positioned on the side walls (Fig. 1d, arrowheads). We transferred gemmalings with nuclei on the side walls to the dark. After 0.5 days of dark treatment, most of the nuclei relocated to the upper walls, and remained there (Fig. 1e). The dark-induced side-to-upper nuclear relocation was inhibited by either the actin-depolymerizing reagent latrunculin B or the microtubule-disrupting reagent oryzalin (Fig. 1f). Simultaneous treatment with both reagents significantly inhibited the relocation (Fig. 1f).

These results indicate that *M. polymorpha* uses both actin and microtubule cytoskeletons for dark-induced side-to-upper nuclear relocation (Fig. 1g). In contrast, in *A. thaliana*, only actin is involved in the dark-induced side-to-bottom nuclear relocation^2,4^.

In *A. thaliana*, nuclear relocation occurs in coordination with chloroplast movement^5^. These movements are driven by two proteins: CHLOROPLAST UNUSUAL POSITIONING 1 (CHUP1)^6^ and KINESIN-LIKE PROTEIN FOR ACTIN-BASED CHLOROPLAST MOVEMENT1 (KAC1)^7^. Phylogenetic analysis confirmed the presence of MpCHUP1 and MpKAC in *M. polymorpha* (Extended Data Fig. 2). We examined two Mp*chup1* mutant alleles (knockout mutant Mp*chup1^KO^*and large deletion mutant Mp*chup1^ld^*) and two Mp*kac* mutant alleles (knockout mutant Mp*kac^KO^* and genome editing mutant Mp*kac^ge^*). None of the mutations in *M. polymorpha* resulted in significant defects in dark-induced nuclear positioning on the upper walls (Fig. 1c and Extended Data Fig. 3), indicating that the relocation is independent of chloroplast movement. This result was supported by imaging data obtained in the dark (Fig. 1a), which show that the distribution of nuclei was different from the distribution of chloroplasts.

In response to intense blue light, vascular plants relocate nuclei to the side walls and keep them there^8^, thereby avoiding ultraviolet B (UVB)-induced DNA damage and cell death^1^. To examine the effect of intense light on the nuclear relocation, we irradiated dark-adapted *M. polymorpha* gemmalings with intense light (300 µmol m^-2^ s^-1^) at various wavelengths (blue, green, red, and far-red) for 3 h. The nuclei relocated from the upper to the side walls specifically in response to blue light (Extended Data Fig. 4a,b). The percentage of nuclei that relocated from the upper to the side walls increased dose-dependently in the range of 50–300 µmol m^-2^ s^-1^ of blue light, but it was not more than 50% (Extended Data Fig. 4c).

Surprisingly, in the presence of prolonged (> 3 h) exposure to intense blue light (300 µmol m^-2^ s^-1^), the nuclei that had been moved to the side walls returned to the upper walls and remained there (Fig. 2a and Extended Data Fig. 4a). The intense light-induced nuclear relocation of *M. polymorpha* therefore consisted of two phases (Fig. 2a): the upper-to-side nuclear relocation within 3 h (Phase I) followed by the side-to-upper nuclear relocation that persisted over a longer period (Phase II). The dual-phase nuclear relocation was entirely correlated with the chloroplast movement, in which chloroplasts were relocated to the side walls and then to the upper walls (Fig. 2b).

**Fig. 2.**
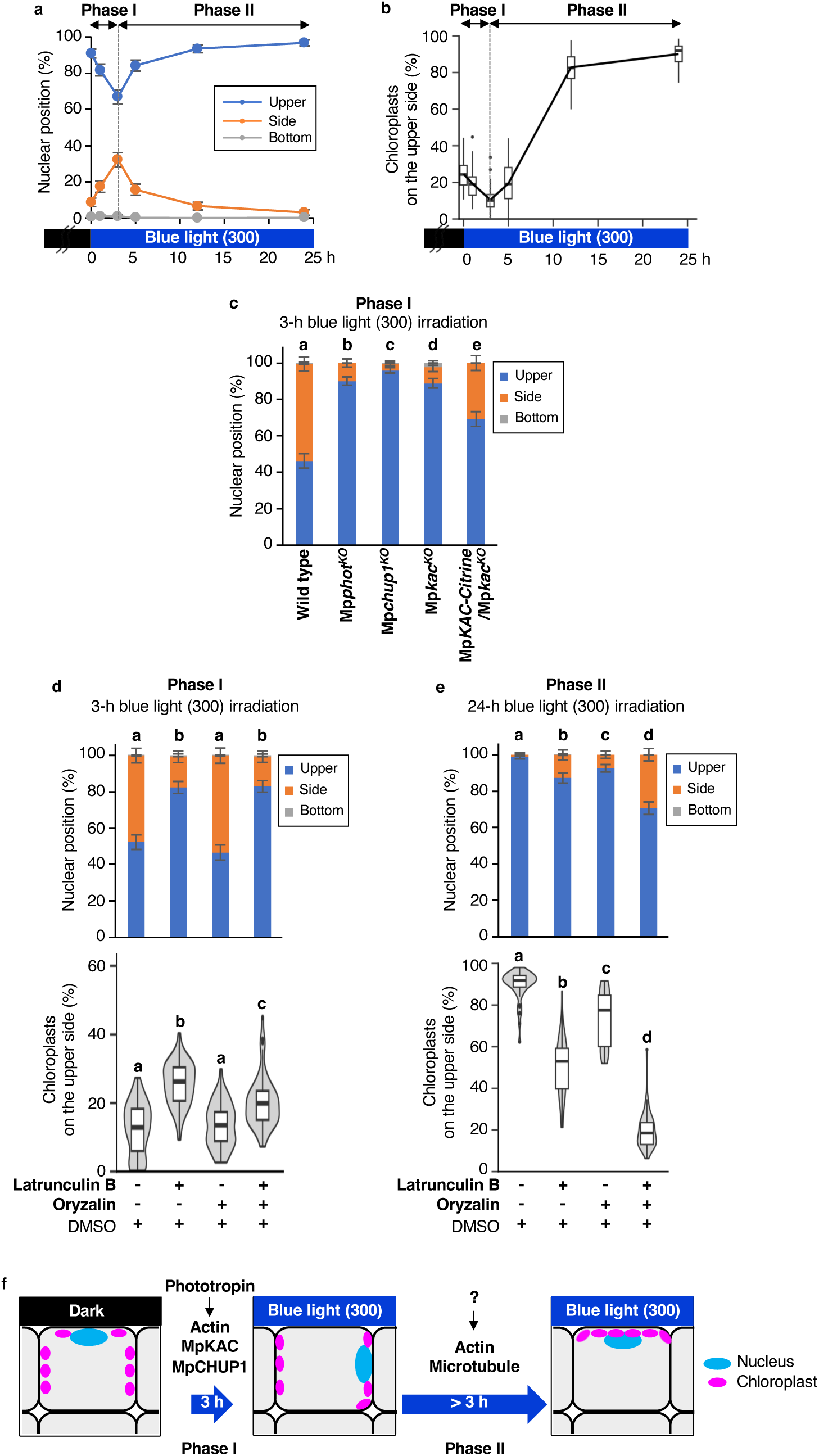
| Intense blue light-induced nuclear relocation consists of two phases under the control of different motile systems in *M. polymorpha*. a, Time course of nuclear relocation under continuous 300 µmol m^-2^ s^-1^ blue light. Ratios of nuclei at each position are shown. Error bars indicate the 95% CI; n = 457–549 cells, five independent experiments. **b,** Time course of chloroplast movement under continuous 300 µmol m^-2^ s^-1^ blue light. Ratio of chloroplasts on the cell surface are shown in the boxplot; n = 50 cells, five independent experiments. **c,** Blue light-induced nuclear relocation in the wild type, Mp*phot^KO^*, Mp*chup1^KO^*, Mp*kac^KO^*, and Mp*KAC*-rescued Mp*kac^KO^*. Dark-adapted gemmalings were irradiated with 300 µmol m^-2^ s^-1^ blue light for 3 h. Ratios of nuclei at each position are shown. Error bars indicate the 95% CI; n = 494–693 cells, five independent experiments. See Extended Data Fig. 4d for Mp*chup1^ld^* and Mp*kac^ge^*. **d,e,** Effects of cytoskeletal inhibitors on blue light-induced nuclear relocation (upper) and chloroplast movement (lower). Dark- adapted gemmalings were irradiated with 300 µmol m^-2^ s^-1^ blue light for 3 h (**d**) and 24 h (**e**). Gemmalings were exposed to inhibitors (either 10 µM latrunculin B, 10 µM oryzalin, or both) starting 1 h before irradiation (**d**) and starting 3 h after irradiation (**e**). Ratios of nuclei at each position are shown (upper). Error bars indicate the 95% CI; n = 510–600 cells (**d**) and n = 546–678 cells (**e**), five independent experiments. Ratios of chloroplasts on the cell surface are shown in the boxplot and violin plots (lower); n = 50 cells, five independent experiments. **f,** Two modes of intense light-induced nuclear relocation in *M. polymorpha* gemmalings. Different letters above the bars indicate statistically significant difference (**c** to **e**).

The nuclear relocation during phase I was remarkably suppressed in all mutants of Mp*phot^KO^*, Mp*chup1^KO^*, Mp*chup1^ld^*, Mp*kac^KO^*, and Mp*kac^ge^* (Fig. 2c and Extended Data Fig. 4d), indicating that Phase I is regulated by the blue-light photoreceptor phototropin (MpPHOT) and two chloroplast movement-related protein homologues (MpCHUP1 and MpKAC). Cytoskeletal drug analysis showed that the actin cytoskeleton, but not the microtubule cytoskeleton, is involved in Phase I movements of nuclei (Fig. 2d, upper) and chloroplasts (Fig. 2d, lower). These Phase I features of *M. polymorpha* are similar to the intense blue light-induced nuclear relocation from the bottom to the side walls in *A. thaliana*^1,3,5,9^. Hence, *M. polymorpha* has acquired an ability to respond to intense blue light, in which nuclei are carried through actin-dependent chloroplast movements that involve MpCHUP1 and MpKAC (Fig. 2f, Phase I).

On the other hand, Phase II is unique to *M. polymorpha* (see Fig. 4a). Phase II involved both the actin and microtubule cytoskeletons in the movements of nuclei (Fig. 2e, upper) and chloroplasts (Fig. 2e, lower), indicating that the nuclear relocation mechanism in Phase II is different from that in Phase I. Prolonged exposure to intense blue light may cause desensitization to phototropin as previously reported^10^, which would decrease the ability of the nuclei to remain on the side walls. In phase II, the nuclei and chloroplasts positioned on the upper walls directly face the blue light (Fig. 2f, Phase II), which causes UV-induced DNA damage. This appears to explain why the sun-loving plant *A. thaliana* has evolved only Phase I^1^. On the other hand, *M. polymorpha*, which is unable to survive in prolonged intense light^11^, thrives in the shade (see Fig. 4b).

Finally, we examined the effect of low light, which has no effect on nuclear relocation in *A. thaliana* (Extended Data Fig. 5). When 3-day-old dark-adapted gemmalings, in which most nuclei were on the upper walls, were exposed to low light (30 µmol m^-2^ s^-1^), their nuclei did not move in a day, but then were gradually relocated to the side walls over several days (Fig. 3a and Extended Data Fig. 4e). The response to low light of *M. polymorpha* was reversible (Fig. 3a). It should be noted that the conditions of continuous low light exposure for days at a time are impossible in nature.

**Fig. 3.**
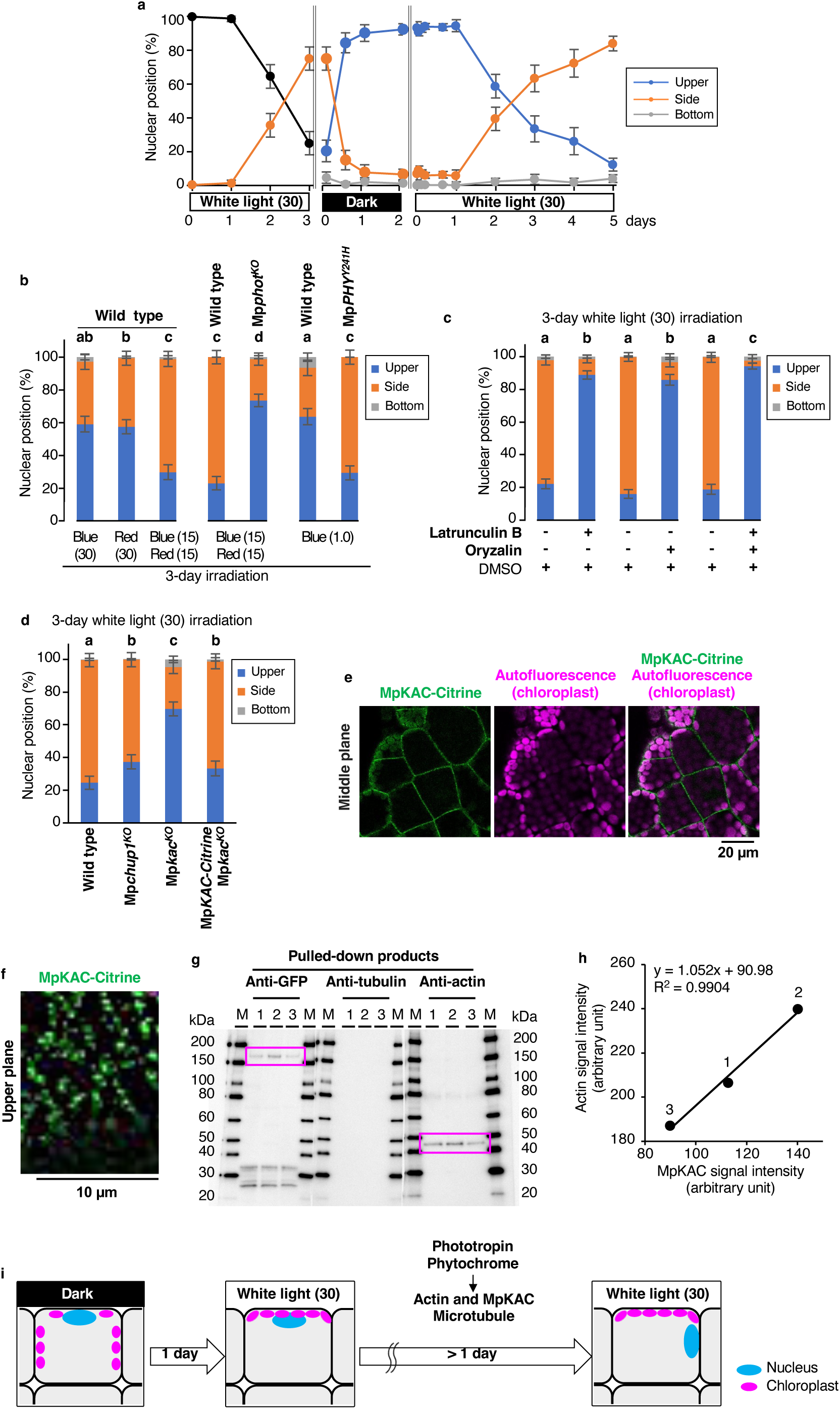
| Low light-induced nuclear relocation requires two photoreceptors and two cytoskeletal components in *M. polymorpha*. a, Time course of nuclear relocation under low light and dark conditions. Gemmalings were grown under continuous 30 µmol m^-2^ s^-1^ white light for 3 days (left), followed by 2 days in the dark (middle) and 5 days under continuous 30 µmol m^-2^ s^-1^ white light (right). Ratios of nuclei at each position are shown. Because gemmae are too small to distinguish nuclei between the upper and bottom wall (left), their nuclei were categorized into two positions (side and the other). Error bars indicate the 95% CI; n = 115–755 cells, five independent experiments. Data are derived from Fig. 1e (middle). **b,** Dependence of nuclear relocation on light wavelength and photoreceptors. Gemmae of the wild type, Mp*phot^KO^*, and *_pro_EF:*Mp*PHY^Y241H^* were irradiated for 3 days with blue and/or red light. Numbers in parentheses represent light intensities (µmol m^-2^ s^-1^). Ratios of nuclei at each position are shown. Error bars indicate the 95% CI; n = 366–511 cells, five independent experiments. **c,** Effects of cytoskeletal inhibitors on low light-induced nuclear relocation. Ratios of nuclei at each position are shown. Error bars indicate the 95% CI; n = 443–737 cells, five independent experiments. **d,** Low light-induced nuclear relocation in the wild-type, Mp*chup1^KO^*, Mp*kac^KO^*, and Mp*KAC*-rescued Mp*kac^KO^*. Ratios of nuclei at each position are shown. Error bars indicate the 95% CI; n = 417–482 cells, five independent experiments. See Extended Data Fig. 4f for Mp*chup1^ld^* and Mp*kac^ge^*. **e,** Distribution of MpKAC-Citrine in the middle plane of the cell of Mp*KAC*-rescued Mp*kac^KO^* gemmaling. MpKAC-Citrine (green); chloroplasts (chlorophyll autofluorescence, magenta). **f,** The magnified image of the upper plane of the cell of Mp*KAC*-rescued Mp*kac^KO^* gemmaling. MpKAC-Citrine (green). **g,** Immunoblots of the anti-GFP antibody pull-down products from three independent transgenic plants expressing MpKAC-Citrine with each of anti-GFP, anti- tubulin, and anti-actin antibodies. M, marker. **h,** Correlation of MpKAC and actin signals in magenta boxes in **g**. **i,** Two modes of low light-induced nuclear relocation in *M. polymorpha* gemmalings. Different letters above the bars indicate statistically significant difference (**b** to **d**).

**Fig. 4.**
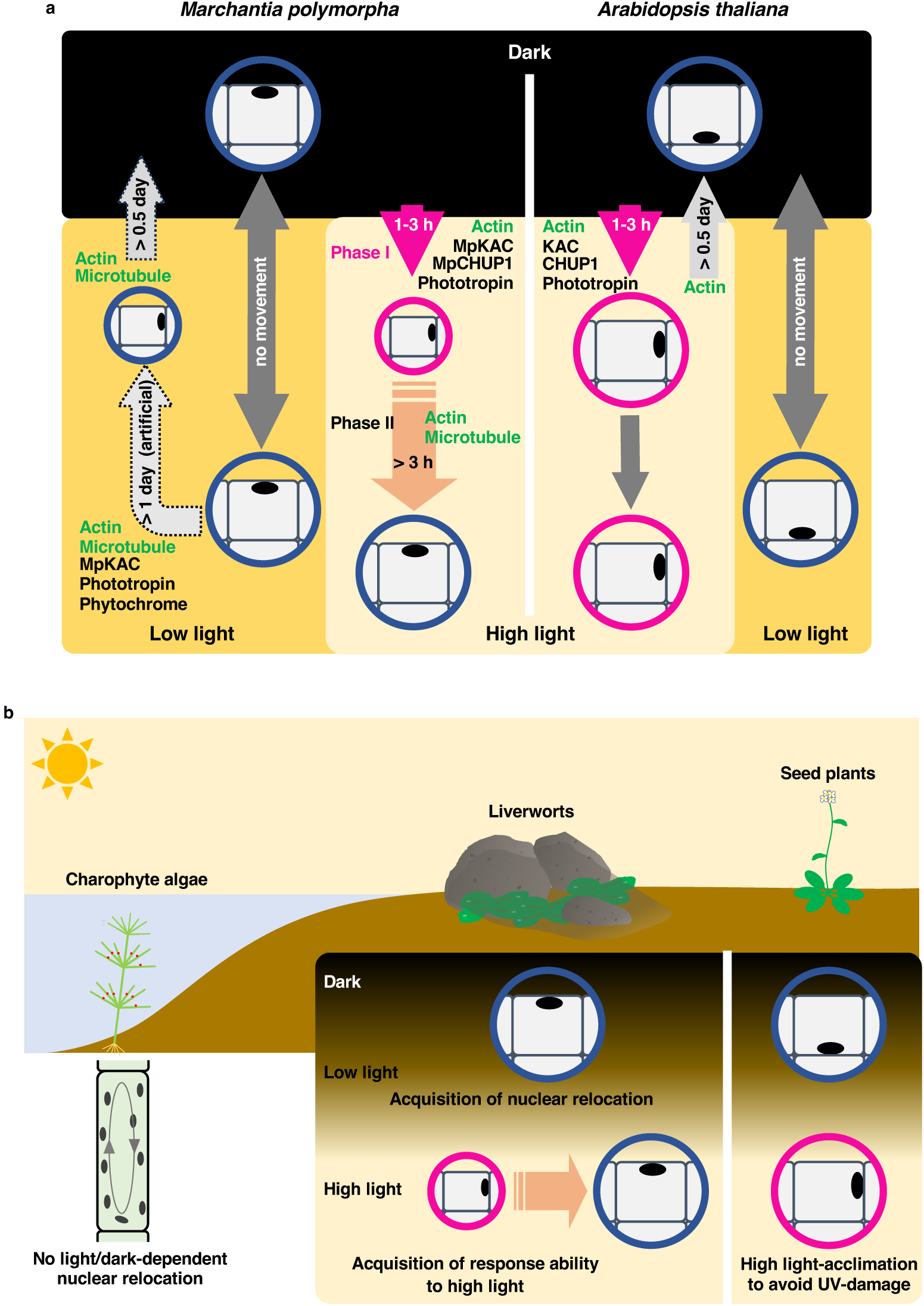
| Evolutionary acquisition and diversification of light- and cytoskeleton-dependent nuclear relocation in land plants. a, Comparison of light- and cytoskeleton-dependent nuclear relocation of *M. polymorpha* (see Fig. 1g, Fig. 2f, and Fig. 3i) with those of *A. thaliana*^2,4^. Note that the phototropin- and actin-dependent nuclear relocation (Phase I) of *M. polymorpha* is similar to that of *A. thaliana* (magenta circles). **b,** Diagram showing our model, in which light-dependent nuclear relocation has initially developed in the common ancestor of land plants as a result of terrestrialization and then diverged as land plants evolved and adapted to new environments. See the text for explanations.

To clarify the mechanism underlying the nuclear relocation unique to *M. polymorpha*, we first investigated the effect of light quality. Weak irradiation of either blue or red light caused around 40% of the nuclei to relocate to the side walls, while their simultaneous irradiation caused more than 70% of the nuclei to relocate to the side walls (Fig. 3b, Wild type). Consistently, deficiency of phototropin (Mp*phot^KO^*) caused a defect in the upper-to-side nuclear relocation (Fig. 3b). The upper-to-side relocation that was induced by very low blue light at 1 µmol m^-2^ s^-1^ was enhanced by expression of a constitutively active form of the red-light photoreceptor phytochrome (Mp*phy^Y241H^*) (Fig. 3b). In addition, the low light-induced upper-to-side nuclear relocation was significantly inhibited by latrunculin B and/or oryzalin (Fig. 3c). Collectively, these results show that the low light-induced nuclear relocation in *M. polymorpha* is regulated coordinately by two photoreceptors (phototropin and phytochrome) and two cytoskeletons (actin and microtubules) (Fig. 3i). This is in contrast to the lack of involvement of microtubules and phytochrome in the nuclear relocation in *A. thaliana*^2,4^.

Neither of the Mp*chup1^KO^* and Mp*chup1^ld^*mutants, which are defective in chloroplast movement, showed a significant defect in nuclear relocation after 3 days of low light exposure (Fig. 3d and Extended Data Fig. 4f), indicating that chloroplast movement contributed minimally to nuclear relocation. This was consistent with the finding that chloroplasts remained on the upper walls in the presence of low light (Fig. 1d). Hence, nuclear positioning in low light is independent of chloroplast movement. On the other hand, the Mp*kac^KO^* and Mp*kac^ge^* mutants showed a significant defect in the low light-induced upper-to-side nuclear relocation (Fig. 3d and Extended Data Fig. 4f). The defect was rescued by expressing Mp*KAC-Citrine* under the control of the native MpKAC promoter (Fig. 3d).

These results indicate that MpKAC plays a substantial role in the response to low light and that Mp*KAC-Citrine* is functional in the cells.

In the rescued line, MpKAC-Citrine was distributed near the plasma membrane (Fig. 3e and Extended Data Fig. 6), as recently reported^12^. Notably, MpKAC-Citrine was also observed as speckles along filamentous structures in the cytosol (Fig. 3f). To clarify the relationship between MpKAC and the cytoskeleton, extracts of the rescued line were subjected to immunoprecipitation with anti-GFP (Citrine) antibody. Three independent experiments showed that the pulled-down products gave positive signals on immunoblots with anti-GFP and anti-actin antibodies, but not with anti-tubulin antibody (Fig. 3g). In the pulled-down products, the signal intensity of MpKAC-Citrine completely paralleled that of actin (Fig. 3h). These results indicate that MpKAC associates with actin but not with tubulin, as does *A. thaliana* KAC1, which functions in the stabilization of actin filaments^7^. Hence, MpKAC regulates low light-induced upper-to-side nuclear relocation by stabilizing actin filaments in *M. polymorpha* (Fig. 3i).

The present results show a clear difference in nuclear positioning in the dark between *M. polymorpha* and *A. thaliana*: *M. polymorpha* positions nuclei on the upper walls of the epidermal cells, whereas *A. thaliana* positions them on the bottom walls (Fig. 4a), resulting in genetic material being kept farther from external environmental stresses. *M. polymorpha*, an early land plant bryophyte with underdeveloped vascular bundles, lives in moist habitats, but such places are also conducive to the growth of pathogens (Fig. 4b). Because the upper walls of the epidermal cells are the outermost parts of the plant body, the nuclei in these cells are necessarily and directly exposed to various environmental stresses including pathogen infection. These cells are able to respond rapidly to such stresses.

Light-dependent nuclear relocation has been observed in a pteridophyte fern^13–15^ and a flowering plant^3^, suggesting that it is a common phenomenon in land plants. Thus, it is of interest to know whether light-dependent nuclear relocation is also present in algae. We checked the nuclear relocation in a charophyte alga (*Chara corallina*), which was reported as the closest living relatives of land plants^16^. The nuclei did not move in response to light (Extended Data Fig. 7). In *Spirogyra crassa* of Zygnematophyceae, which was reported as the most likely sister group of land plants^17^, the nucleus seems to be kept in the center of the cell^18^. These results suggest that light-dependent nuclear relocation originated in the common ancestor of land plants. Thus, our results provide a conceptual advance in how light-dependent nuclear relocation was acquired in land plants (Fig. 4b): it appears to have initially developed as a result of terrestrialization and then diverged as land plants evolved and adapted to new environments.

## Methods

### Plant materials and growth conditions

A male *Marchantia polymorpha* Takaragaike-1 (Tak-1) was used as a wild-type plant. The phototropin mutant (*Mpphot^KO^*)^19^, constitutively active mutant for phytochrome (*_pro_EF:MpPHY^Y241H^*)^20^, Mp*chup1*-knockout mutant (Mp*chup1^KO^*)^12^, Mp*chup1* large-deletion mutant (Mp*chup1^ld^*)^12^, Mp*KAC*-knockout mutant (Mp*kac^KO^*)^12^, Mp*KAC* genome-edited mutant (Mp*kac^ge^*), and MpKAC-Citrine-expressing plants had the Tak-1 background. The plants were maintained asexually as previously described^21,22^. Gemmae were grown aseptically on germination medium (one-half-strength Gamborg’s B5 salts, 0.025% [w/v] MES-KOH, pH 5.7, and 1% [w/v] agar) at 22 °C under continuous 30 µmol m^-2^ s^-1^ white light conditions, and the 3-day-old gemmalings (the early growth stage of thalli developing from gemmae) were used unless otherwise stated. *Arabidopsis thaliana* (ecotype Columbia, *gl1* background) was grown at 22 °C under a 8-h-white light (35 µmol m^-2^ s^-1^)/16-h-dark cycle.

*Chara corallina* was grown at 25 °C under a 12-h-white light (20 µmol m^-2^ s^-1^)/12-h-dark cycle. The light intensity was measured using a quantum sensor (LI-190SA; LI-COR) or light analyzer (LA-105; NK System).

### Generation of MpKAC-Citrine expressing plants

We constructed Mp*KAC-Citrine* for Mp*kac^ko^* complementation through amplification of the genomic region containing the 2880-bp upstream of the translational initiation site and the coding sequence of Mp*KAC* from Tak-1 gDNA using the primers 5′- AACCAATTCAGTCGACCAAGCCCAGGTGTATATGGA-3′ and 5′- AAGCTGGGTCTAGATATCCATGAGGCTCATCTCCACCAA-3′, subcloned it into pENTR1A between the SalI and EcoRV sites using an In-Fusion HD Cloning Kit (Takara), and transferred it to the destination vector pMpGWB307^23^ using Gateway Clonase II Enzyme Mix (Thermo Fisher Scientific). The resulting plasmids were introduced into Mp*kac^ko^* using the G-AgarTrap method^24^.

To construct *_pro_*Mp*EF1α:*Mp*KAC-Citrine*, we amplified the coding sequence of Mp*KAC* using the primers 5′-AACCAATTCAGTCGACATGGCAGAGAGGAATAGCTGG-3′ and 5′- AAGCTGGGTCTAGATATCCATGAGGCTCATCTCCACCAA-3′, subcloned it into pENTR1A between the SalI and EcoRV sites using the In-Fusion HD Cloning Kit (Takara), and transferred it to the destination vector pMpGWB108. The resulting plasmids were introduced into Tak-1 cells using the G-AgarTrap method^24^.

### Genome editing of Mp*KAC*

To generate Mp*kac^ge^* mutants, we edited the Mp*KAC* locus using a CRISPR/Cas9-based genome-editing system as previously described^25^. pMpGE013 was digested with AarI restriction enzyme. Annealed oligos for MpKAC_gRNA1_f (5′- CTCGTCAAAATTACGCGAGCCACT-3′) and MpKAC_gRNA1_r (5′-AAACAGTGGCTCGCGTAATTTTGA-3′) were ligated into the linearized vector using DNA Ligation Kit Mighty Mix (Takara). The resulting plasmids were introduced into Tak-1 using the G-AgarTrap method^24^. We extracted gDNA from T_1_ thalli and analyzed the sequences using PCR and direct sequencing to identify frameshift mutations.

### Dark and light treatments

The dark treatment of *M. polymorpha* gemmalings occurred over 2 days unless otherwise stated. *C. corallina* was dark-treated for 1 day. For the analysis of light-induced nuclear relocation, the samples were irradiated with blue (470 nm), green (525 nm), red (660 nm), or far-red (735 nm) light using an LED light source system (IS-mini; CCS). The light intensity was measured using the quantum sensor (LI-190SA; LI-COR) or light analyzer (LA-105; NK System).

### Visualization of the nucleus, chloroplast, and MpKAC-Citrine

Epidermal cells near the apical notches of *M. polymorpha* gemmalings, adaxial pavement and palisade mesophyll cells of *A. thaliana* leaves, and branchlet cells of *C. corallina* were observed using a fluorescence microscope (Axio Scope A1; Zeiss) equipped with a CCD camera (Axiocam 506 color; Zeiss) or confocal microscope (LSM800; Zeiss: SP8; Leica). To visualize the nuclei, the samples were fixed in a buffer solution (50 mM PIPES, 10 mM EGTA, and 5 mM MgSO_4_, pH 7.0) containing 1% (w/v) glutaraldehyde for 1 h, stained with 1 µg/mL DAPI (Sigma-Aldrich), and diluted in fixation buffer supplemented with 0.03% (v/v) Triton X-100 for 1.5 h. Chloroplasts were visualized with chlorophyll autofluorescence. MpKAC-Citrine was observed without cell fixation.

### Inhibitor treatments

Stock solutions [5 mM latrunculin B (actin-depolymerizing agent, Abcam) and 50 mM oryzalin (microtubule-disrupting agent, Sigma-Aldrich] were prepared in dimethyl sulfoxide (DMSO) and diluted to 10 µM with distilled water. To analyze dark-induced nuclear relocation, inhibitors were applied to 3-day-old gemmalings for 12 h in the dark. As controls, 0.2% (v/v) DMSO and 0.02% (v/v) DMSO were used for latrunculin B and oryzalin, respectively. To analyze intense light-induced nuclear relocation, dark-adapted gemmalings were irradiated with 300 µmol m^-2^ s^-1^ blue light for 3 h. Gemmalings were exposed to inhibitors (either 10 µM latrunculin B, 10 µM oryzalin, or both) starting 1 h before irradiation. Alternatively, dark-adapted gemmalings were irradiated with 300 µmol m^-2^ s^-1^ blue light for 24 h. Gemmalings were exposed to inhibitors (either 10 µM latrunculin B, 10 µM oryzalin, or both) starting 3 h after irradiation. As a control, 0.22% (v/v) DMSO was used. To analyze low light-induced nuclear relocation, gemmae were grown on germination plates supplemented with inhibitors (10 µM) under 30 µmol m^-2^ s^-1^ white light conditions for 3 days. As a control, gemmalings were grown on germination plates supplemented with 0.2% (v/v) DMSO for latrunculin B or 0.02% (v/v) DMSO for oryzalin.

### Pull-down assay

A pull-down assay was performed using green fluorescent protein (GFP)-Trap agarose (ChromoTek). We homogenized 1 g of 24-d-old MpKAC-Citrine overexpressing plants in buffer containing 137 mM NaCl, 2.7 mM KCl, 10 mM Na_2_HPO_4_•12H_2_O, 1.8 mM KH_2_PO_4_, 0.5 mM EDTA, 0.1% (v/v) Triton X-100, protease inhibitor cocktail (Roche), and 2.5 mM DSP (pH 7.4) and incubated for 30 min at 25 °C. The homogenates were centrifuged at 9100 ×*g* and 4 °C to remove cellular debris. The supernatants were mixed with GFP-Trap Agarose and incubated at 4 °C for 1 h with inverted mixing. After washing the GFP-Trap Agarose three times with buffer containing 50 mM Tris-HCl, 150 mM NaCl, 0.5 mM EDTA, 0.1% (v/v) Triton X-100, and a protein inhibitor cocktail (pH 7.5), immunoaffinity complexes were eluted with sample buffer containing DTT and SDS (Atto).

### SDS-PAGE and immunoblot analysis

Protein extracts obtained from the pull-down assay were subjected to SDS-PAGE, followed by immunoblot analysis, as previously described^4^. SDS-PAGE was performed using a conventional 4–15% precast gel (Criterion TGX; Bio-Rad) according to the manufacturer’s instructions. Immunoreactive signals were detected using an ECL detection system (GR Healthcare) with anti-GFP antibody (1:5,000; JL-8; Clontech), anti-actin antibody (1:2,000; clone 10-B3; Sigma-Aldrich), anti-microtubule antibody (1:2,000; clone B-5-1-2; Sigma-Aldrich), and ECL anti-mouse IgG horseradish peroxidase-linked whole antibody (1:1,000; GE Healthcare).

### Measurement of chloroplasts and cell surface

The areas of the chloroplasts and cell surface (upper cell plane) were measured using Fiji software (version: 2.14.0/1.54h; https://imagej.net/software/fiji/)^26^ to evaluate chloroplast movement. Cells of interest were traced on bright-field images, and their outlines were generated at 15 pixels smaller than the original outlines. The region enclosed by the small outline was defined as the upper surface of the cell, and the number of pixels was counted. Chlorophyll autofluorescence images were binarized using Fiji software, the smaller outlines were adapted to the binarized images, and the number of pixels of chlorophyll autofluorescence within the small outlines was determined. We finally determined the ratio of chlorophyll fluorescence to cell-surface pixels.

### Phylogenetic analysis

*CHUP1* and *KAC*, as well as their outgroup genes, in *A. thaliana* (At), *Oryza sativa* (Os), *M. polymorpha* (Mp), *Physcomitrella patens* (Pp)^27,28^, and *Chara braunii* (Cb) were searched with AtCHUP1 and AtKAC1 using BLASTp on the NCBI website (https://www.ncbi.nlm.nih.gov/). The accession numbers used in phylogenetic analysis were shown in Supplementary Table 1. The gene accession numbers for the Mp genes (Supplementary Table 1) were obtained from MarpolBase (https://marchantia.info/). Multiple alignments were prepared using MAFFT v7.490^29,30^, and non-homologous regions were trimmed using trimAl v1.4^31^ with the “-gappyout” option. A phylogenetic tree was constructed using the maximum likelihood method with RAxML v8.2.12^32^, with 1,000 bootstraps. The evolutionary model parameter was estimated using the “-m PROTGAMMAAUTO” option. The phylogenetic tree was visualized using FigTree v1.4.4 (http://tree.bio.ed.ac.uk/software/figtree/).

### Statistical analysis

Statistical analysis and creation plots were done by Microsoft Excel (Microsoft Corporation) and R version 4.1.2 (https://www.R-project.org/)^33^ with Rstudio 2024.04.2+764 (http://www.rstudio.com/)^34^. The ratios of nuclear positions were calculated from all cells observed from five individuals in each experiment. The data were plotted as stacked bar graphs or line graphs with Microsoft Excel and the error bars showed 95% confidence interval approximating Gaussian distribution. Comparison between experimental groups were tested by Fisher’s exact test with Bonferroni correction. The ratios of chloroplast positions were calculated as follows; for each experimental condition, 10 cells were observed from each individual, for a total of five individuals with 50 cells, and the ratio of chloroplast position was calculated for each cell. The 50 ratio-data obtained for each experiment were plotted as box and violin or line plots with R package ggplot2^35^. Comparison between experimental groups were tested by Mann-Whitney *U*-test with Bonferroni correction. All tests were two tailed, and statistical significance was set at *p* < 0.05.

## Data availability

Sequence data from this study can be found in MarpolBase data libraries under the following accession numbers: Mp*PHOT* (Mp5g03810), Mp*PHY* (Mp2g16090), Mp*CHUP1* (Mp7g03540), and Mp*KAC* (Mp3g03760). The data that support the findings of this study are available from the corresponding authors upon request.

## Acknowledgments

We are deeply grateful to Shingo Takagi (Osaka University, deceased on October 28, 2023) for his invaluable suggestions for this study. We are grateful to Yoshiji Okazaki (Osaka Medical College) for the donation of *Chara corallina*. We thank James Raymond (Eigoken; University of Nevada, Las Vegas) for his critical reading of the manuscript. This work was supported by a Grant-in-Aid for Specially Promoted Research (no. 22000014 to I.H.-N.), by Grants-in-Aid for Scientific Research (no. 15H05776 to I.H-.N., no. 17K15145 to K.I., and no. 18H05496 to H.U.) from the Japan Society for the Promotion of Science (JSPS), and by the Hirao Taro Foundation of KONAN GAKUEN for Academic Research to I.H.-N.

## Author contributions

Conceptualization, Project administration, and Writing; K.I. and I.H.-N.: Supervision; K.I., H.U., and I.H.-N.: Investigation and Data curation; K.I.: Formal analysis; K.I. and H.Y.: Visualization; K.I., H.Y., and I.H.-N.: Methodology; Y.W.: Resources; K.C.M., A.K., Y.S.,

T.S., R.N., T.K., and A.H.: Funding acquisition; K.I., A.H., H.U., and I.H.-N.

## Competing interests

The authors declare no competing interests.

**Correspondence and requests for materials** should be addressed to Kosei Iwabuchi or Ikuko Hara-Nishimura.

---

**Extended Data Fig. 1.**
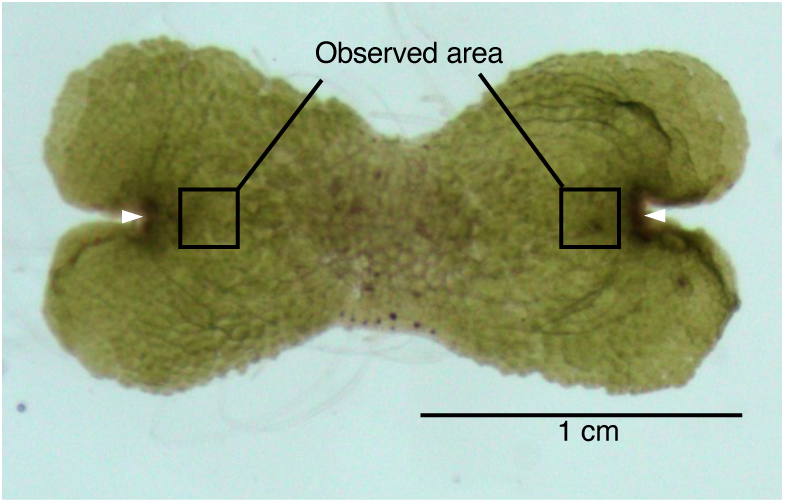
| *M. polymorpha* gemmaling. A 3-day-old wild-type Tak-1 gemmaling. Epidermal cells near the two apical notches (arrowheads) were observed.

**Extended Data Fig. 2.**
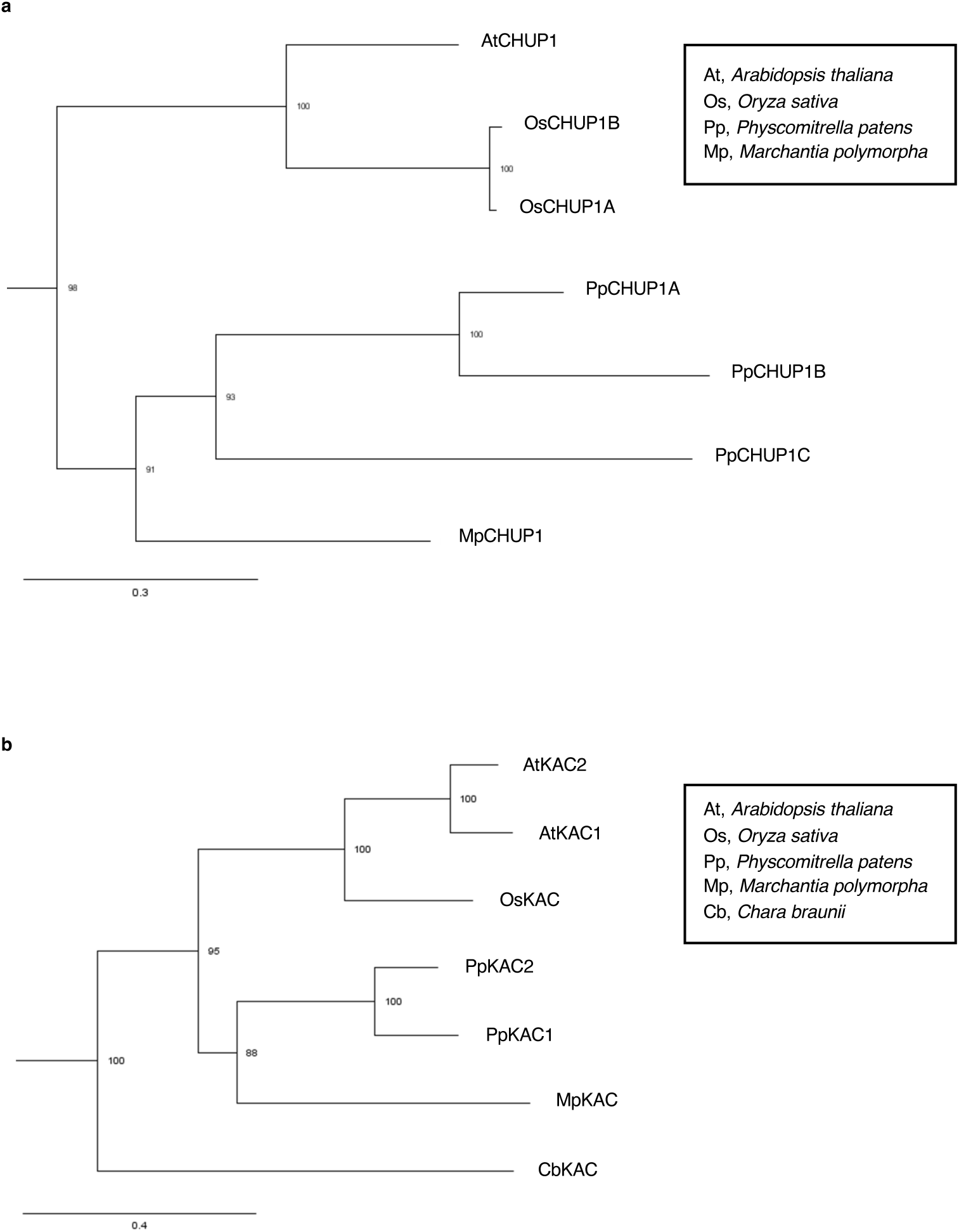
| Phylogenetic trees of CHUP1 and KAC proteins. a,b, Maximum likelihood phylogenetic trees of CHUP1 (**a**) and KAC (**b**) proteins. The numbers near the nodes are the results of the bootstrap analysis (1,000 replicates). Outgroups are not shown.

**Extended Data Fig. 3.**
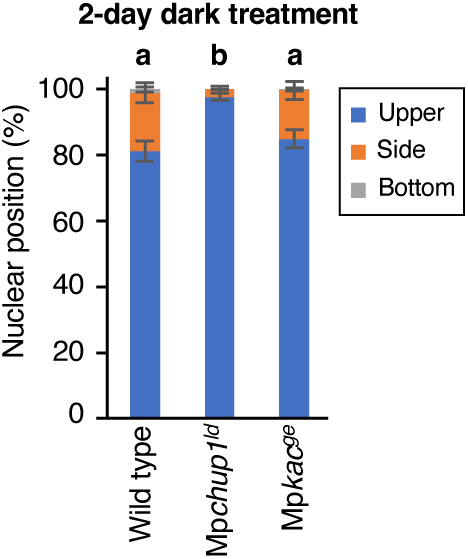
| Dark-induced nuclear relocation in the wild type, Mp*chup1^ld^*, and Mp*kac^ge^*. Ratios of nuclei at each position are shown. Error bars indicate the 95% CI; n = 613–773 cells, five independent experiments. See Fig. 1c for Mp*chup1^KO^* and Mp*kac^KO^*.

**Extended Data Fig. 4.**
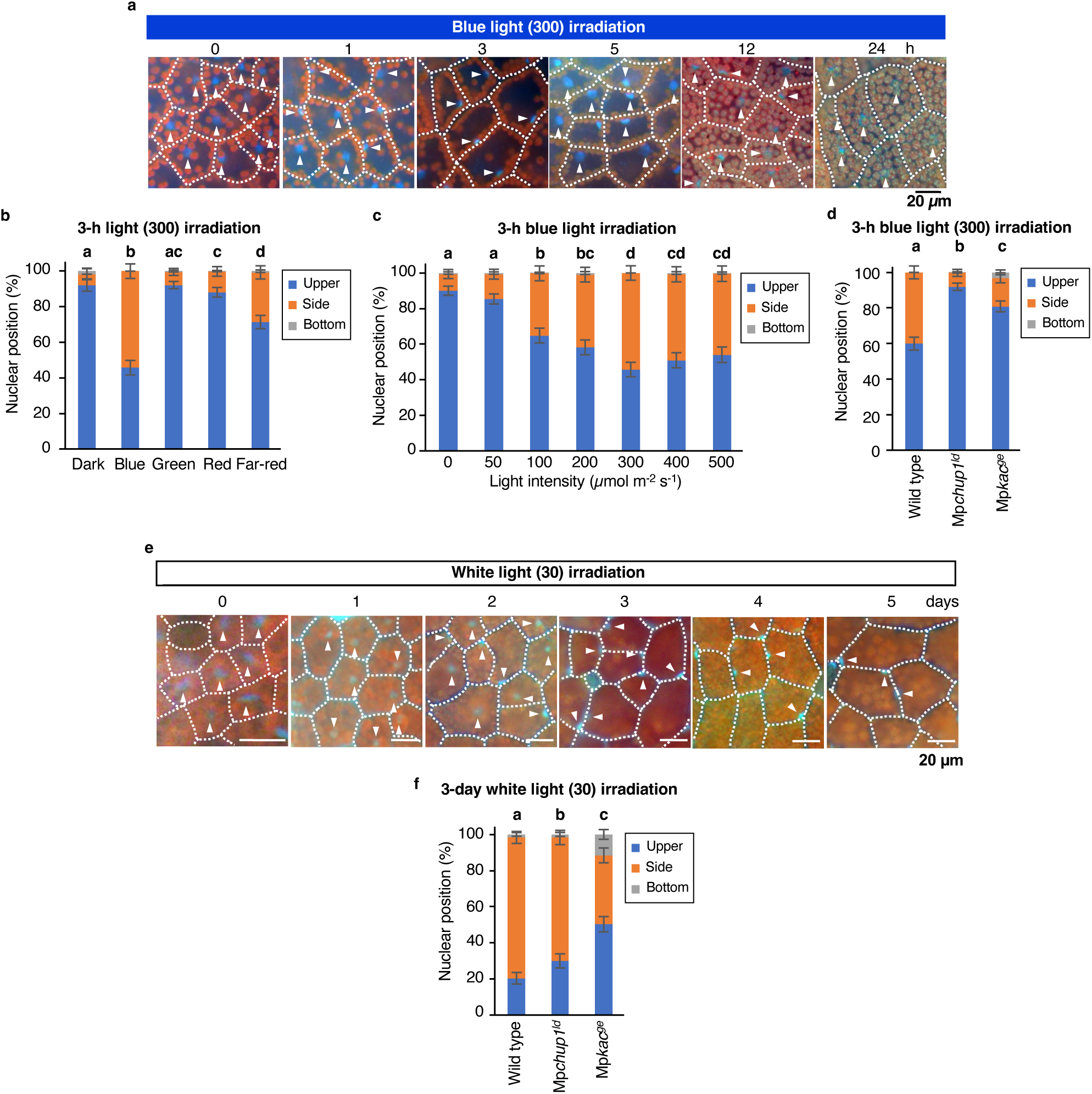
| Intense blue light- and low light-induced nuclear relocation in *M. polymorpha*. a, Dark-adapted gemmalings were irradiated with 300 µmol m^-2^ s^-1^ blue light for 24 h. Representative images of nuclei stained with DAPI (blue, arrowheads) and chloroplasts (chlorophyll autofluorescence, red). Cells are surrounded by dashed lines. **b,** Dependence of nuclear relocation on light wavelength. Dark-adapted gemmalings were irradiated with blue, green, red, or far-red light at 300 µmol m^-2^ s^-1^ for 3 h. Ratios of nuclei at each position are shown. Error bars indicate the 95% CI; n = 230–687 cells, five independent experiments. **c,** Dependence of nuclear relocation on blue light intensities. Dark-adapted gemmalings were irradiated with blue light at 0, 50, 100, 200, 300, 400, and 500 µmol m^-2^ s^-1^ for 3 h. Ratios of nuclei at each position are shown. Error bars indicate the 95% CI; n = 503–602 cells, five independent experiments. **d,** Blue light-induced nuclear relocation in the wild type, Mp*chup1^ld^*, and Mp*kac^ge^*. Dark-adapted gemmalings were irradiated with 300 µmol m^-2^ s^-1^ blue light for 3 h. Ratios of nuclei at each position are shown. Error bars indicate the 95% CI; n = 648–705 cells, five independent experiments. See Fig. 2c for Mp*chup1^KO^* and Mp*kac^KO^*. **e,** Gemmae were irradiated with 30 µmol m^-2^ s^-1^ white light for 5 days. Representative images of nuclei stained with DAPI (blue, arrowheads) and chloroplasts (chlorophyll autofluorescence, red). Cells are surrounded by dashed lines. **f,** Low light-induced nuclear relocation in the wild type, Mp*chup1^ld^*, and Mp*kac^ge^*. Ratios of nuclei at each position are shown. Error bars indicate the 95% CI; n = 533–603 cells, five independent experiments. See Fig. 3d for Mp*chup1^KO^* and Mp*kac^KO^*.

**Extended Data Fig. 5.**
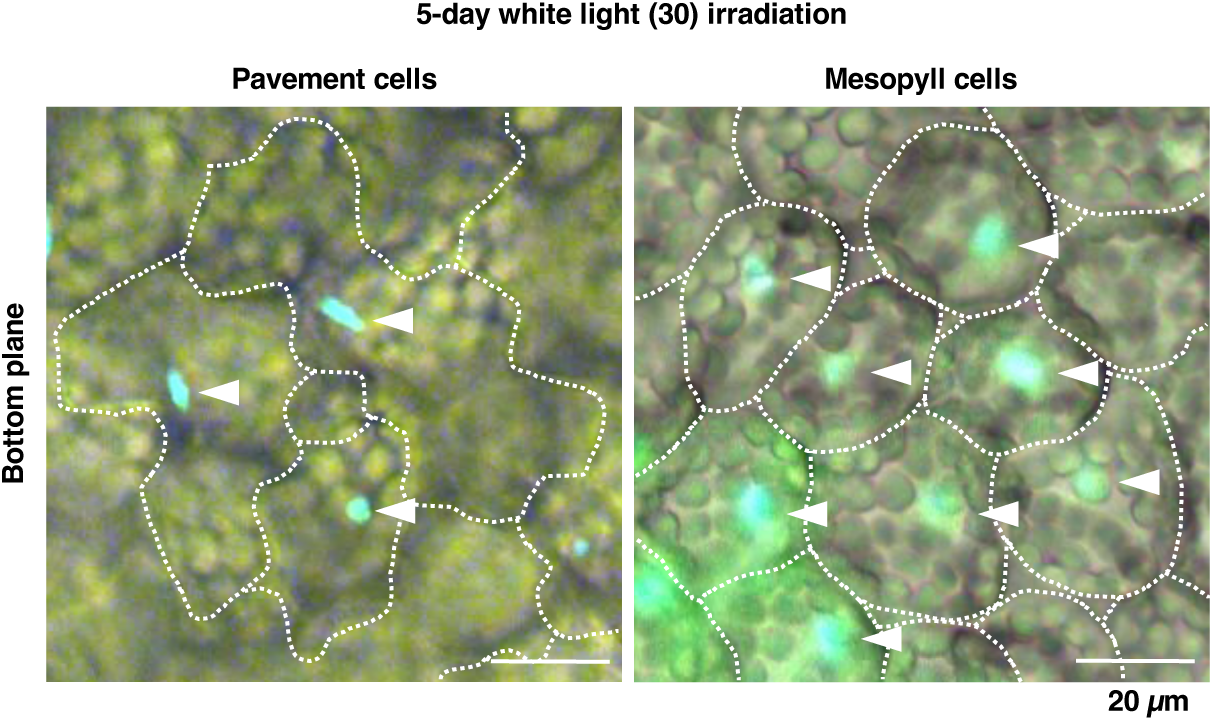
| Distribution of nuclei in *A. thaliana* leaves under prolonged exposure to low light. Representative images of nuclei stained with DAPI (blue, arrowheads) in pavement (left) and mesophyll (right) cells of detached leaves exposed to 30 µmol m^-2^ s^-1^ white light for 5 days. Cells are surrounded by dashed lines.

**Extended Data Fig. 6.**
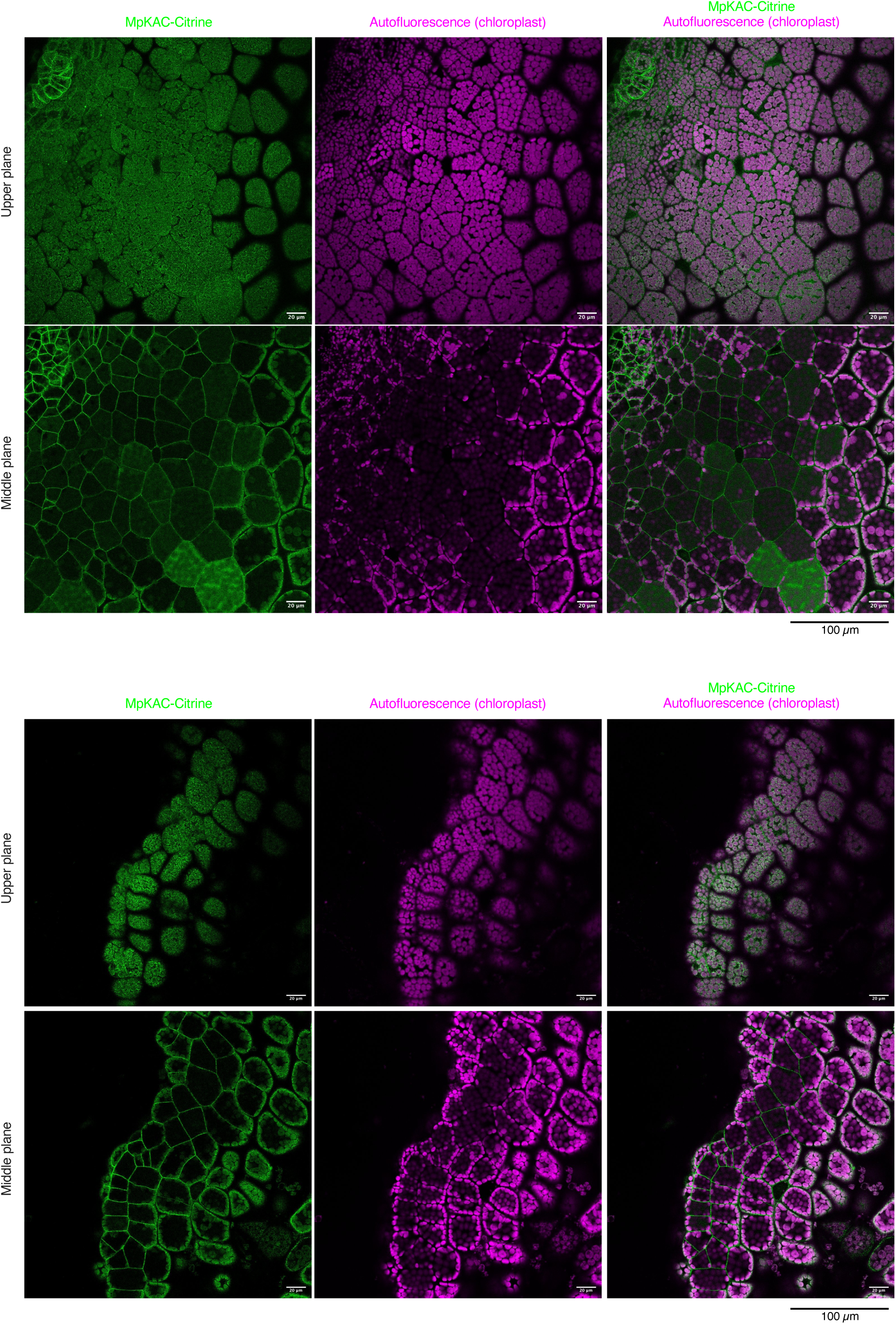
| Distribution of MpKAC-Citrine in the upper and middle planes of Mp*KAC*-rescued Mp*kac^KO^* gemmalings. MpKAC-Citrine (green); chloroplasts (chlorophyll autofluorescence, magenta). Two gemmalings are shown.

**Extended Data Fig. 7.**
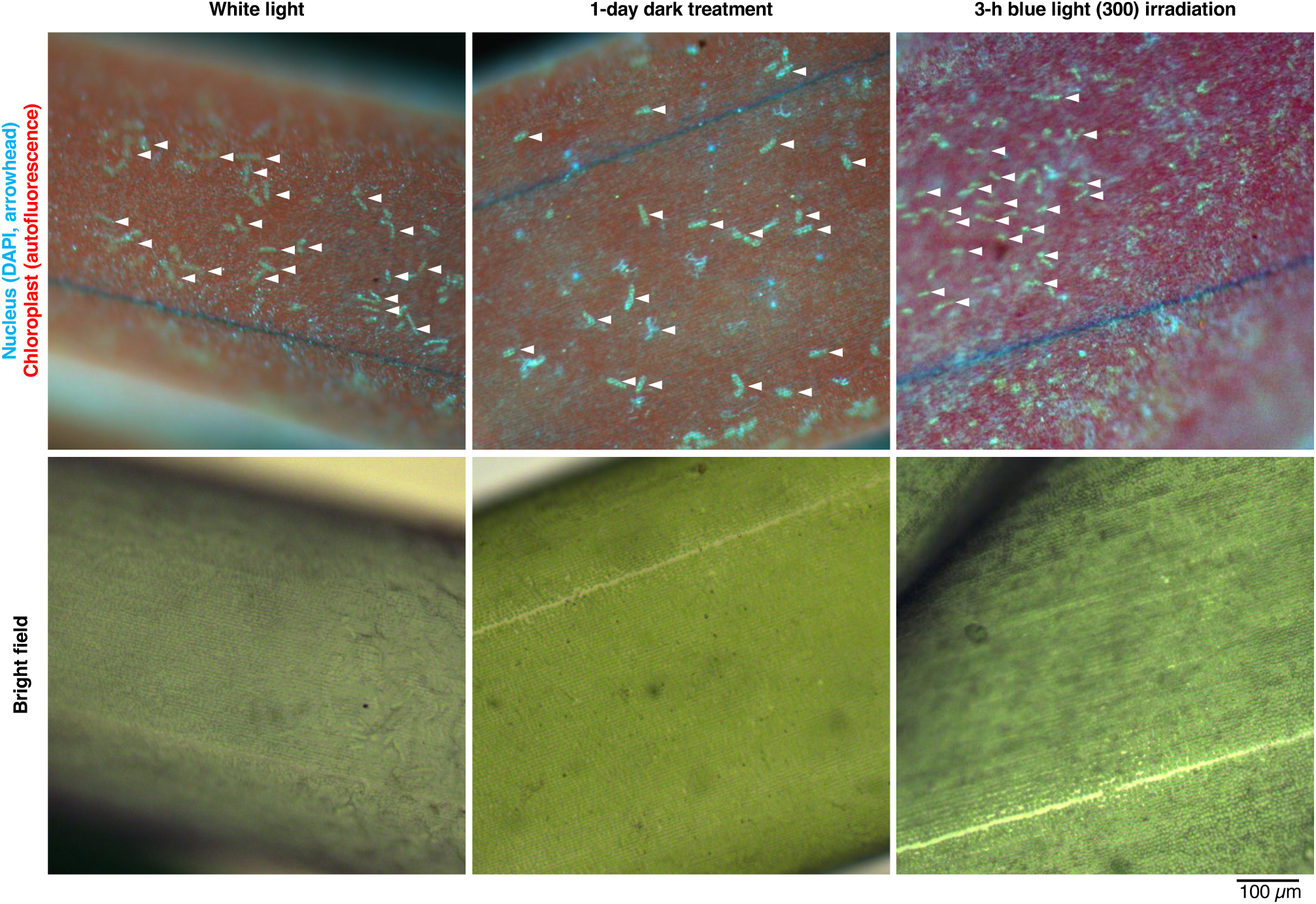
| Distribution of nuclei in a charophyte alga (*C. corallina*) under the dark and light conditions. Nuclei stained with DAPI (blue, arrowhead) and chloroplasts (chlorophyll autofluorescence, red) together with their corresponding bright field images are shown. The branchlet cells of *C. corallina* grown under white light (left) were dark-treated for 1 day (middle) and then irradiated with 300 µmol m^-2^ s^-1^ blue light for 3 h (right).

**Supplementary Table 1.**
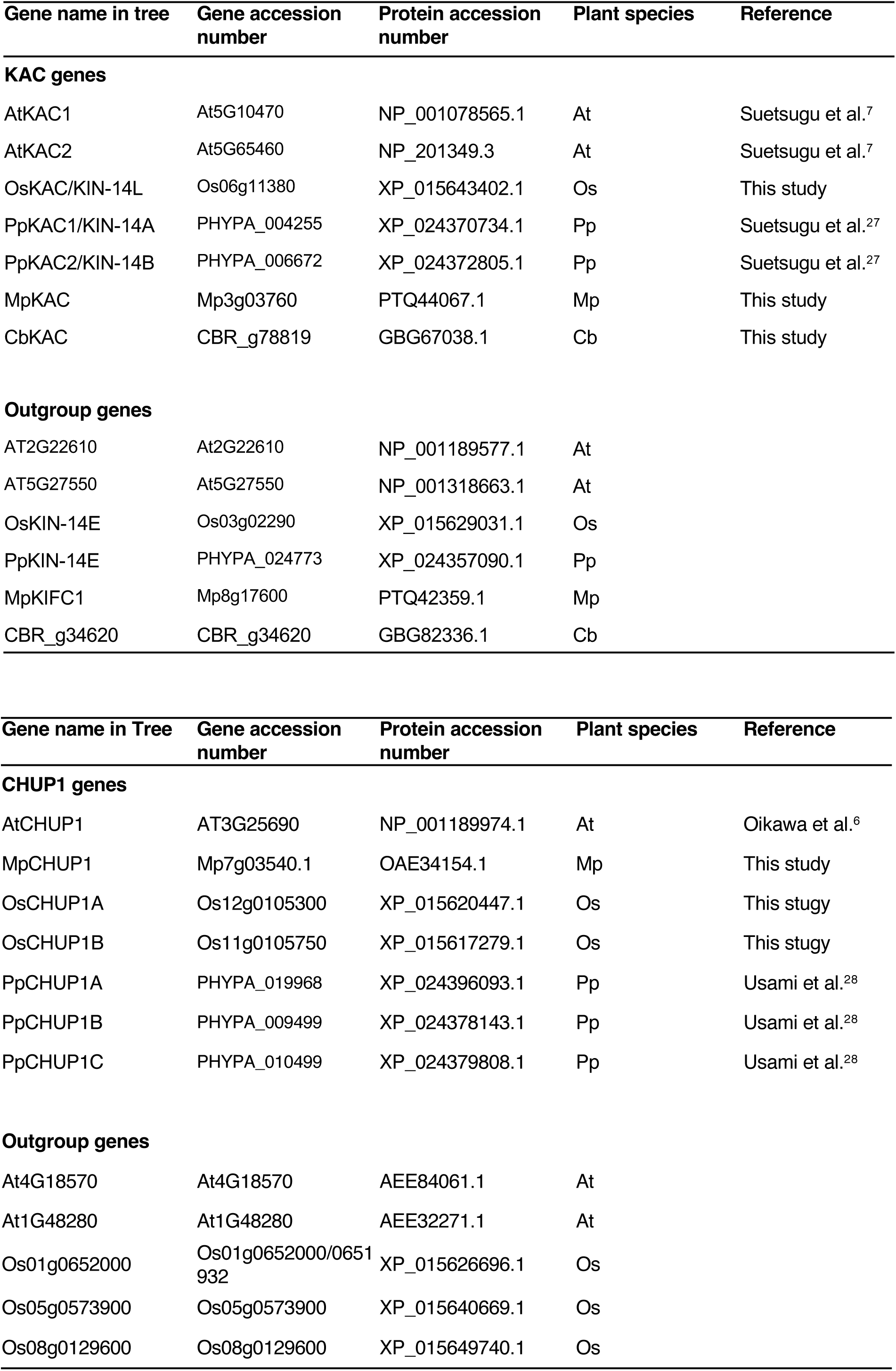
| Accession numbers for phylogenetic analysis.

| Gene name in tree | Gene accession number | Protein accession number | Plant species | Reference |
| --- | --- | --- | --- | --- |
| **KAC genes** |  |  |  |  |
| AtKAC1 | At5G10470 | NP_001078565.1 | At | Suetsugu et al.<sup>7</sup> |
| AtKAC2 | At5G65460 | NP_201349.3 | At | Suetsugu et al.<sup>7</sup> |
| OsKAC/KIN-14L | Os06g11380 | XP_015643402.1 | Os | This study |
| PpKAC1/KIN-14A | PHYPA_004255 | XP_024370734.1 | Pp | Suetsugu et al.<sup>27</sup> |
| PpKAC2/KIN-14B | PHYPA_006672 | XP_024372805.1 | Pp | Suetsugu et al.<sup>27</sup> |
| MpKAC | Mp3g03760 | PTQ44067.1 | Mp | This study |
| CbKAC | CBR_g78819 | GBG67038.1 | Cb | This study |
| **Outgroup genes** |  |  |  |  |
| AT2G22610 | At2G22610 | NP_001189577.1 | At |  |
| AT5G27550 | At5G27550 | NP_001318663.1 | At |  |
| OsKIN-14E | Os03g02290 | XP_015629031.1 | Os |  |
| PpKIN-14E | PHYPA_024773 | XP_024357090.1 | Pp |  |
| MpKIFC1 | Mp8g17600 | PTQ42359.1 | Mp |  |
| CBR_g34620 | CBR_g34620 | GBG82336.1 | Cb |  |

| Gene name in Tree | Gene accession number | Protein accession number | Plant species | Reference |
| --- | --- | --- | --- | --- |
| **CHUP1 genes** |  |  |  |  |
| AtCHUP1 | AT3G25690 | NP_001189974.1 | At | Oikawa et al.<sup>6</sup> |
| MpCHUP1 | Mp7g03540.1 | OAE34154.1 | Mp | This study |
| OsCHUP1A | Os12g0105300 | XP_015620447.1 | Os | This study |
| OsCHUP1B | Os11g0105750 | XP_015617279.1 | Os | This study |
| PpCHUP1A | PHYPA_019968 | XP_024396093.1 | Pp | Usami et al.<sup>28</sup> |
| PpCHUP1B | PHYPA_009499 | XP_024378143.1 | Pp | Usami et al.<sup>28</sup> |
| PpCHUP1C | PHYPA_010499 | XP_024379808.1 | Pp | Usami et al.<sup>28</sup> |
| **Outgroup genes** |  |  |  |  |
| At4G18570 | At4G18570 | AEE84061.1 | At |  |
| At1G48280 | At1G48280 | AEE32271.1 | At |  |
| Os01g0652000 | Os01g0652000/0651932 | XP_015626696.1 | Os |  |
| Os05g0573900 | Os05g0573900 | XP_015640669.1 | Os |  |
| Os08g0129600 | Os08g0129600 | XP_015649740.1 | Os |  |

